# Decoding of attentional state using high-frequency local field potential is as accurate as using spikes

**DOI:** 10.1101/2020.08.31.275347

**Authors:** Surya S Prakash, Aritra Das, Sidrat Tasawoor Kanth, J. Patrick Mayo, Supratim Ray

## Abstract

Local field potentials (LFPs) in visual cortex are reliably modulated when the subject’s focus of attention is cued into versus out of the receptive field of the recorded sites, similar to modulation of spiking activity. However, human psychophysics studies have used an additional attention condition, neutral cueing, for decades. The effect of neutral cueing on spiking responses was examined recently and found to be intermediate between cued and uncued conditions. However, whether LFPs are also precise enough to represent graded states of attention is unknown. We found that LFPs during neutral cueing were intermediate between cued and uncued conditions, and, for a single electrode, attention was more discriminable using high frequency (>30 Hz) LFP power than spikes. Surprisingly, spikes did not outperform LFPs even when discriminability was computed using multiple electrodes. These results constrain the spatial scale attention operates over and highlight the usefulness of LFPs in studying attention.

## Introduction

Power at different frequency bands like alpha (8-12 Hz) and gamma (30-80 Hz) of the local field potential (LFP) is reliably modulated by selective attention (Fries et al., 2001, 2008; Chalk et al., 2010; Khayat et al., 2010; Khamechian et al., 2019) and can be used to predict behavior (Womelsdorf et al., 2006). These results have led to suggestions that gamma oscillations could play a role in routing information across cortical areas (Fries, 2015). However, most of these studies employed a standard attention task in which the animal’s attention was either cued inside (attend-in) or outside (attend-out) the receptive field (RF) of the recording sites.

Additional attentional conditions are often used in psychophysical tasks in humans. Here, a spatial attention task typically involves cueing the subject to deploy attention to a desired location at which a target is anticipated (Posner, 1980). The target usually occurs at the cued location (which are called “valid” trials) but can infrequently also occur at an uncued location (“invalid” trials), such that the behavioral effect of spatial attention can be measured by comparing the performance (in terms of detection rates) and reaction times for valid versus invalid trials (Posner, 1980; Carrasco, 2011). In addition, such studies often use an ambiguous or “neutral” cueing condition where the target change is equally likely at both locations. These studies have shown that behavioral performance during neutral cueing is intermediate between valid and invalid conditions (Posner et al., 1978; Posner, 1980; Mangun and Hillyard, 1990; Montagna et al., 2009). While neurophysiological studies of spatial attention have compared neuronal responses for attend-in versus attend-out conditions for decades (Moran and Desimone, 1985; Treue and Maunsell, 1996; McAdams and Maunsell, 1999), the effect of neutral cueing on spiking activity has been studied only recently, and found to be intermediate between cued and uncued conditions (Mayo and Maunsell, 2016) (but see (Denfield et al., 2018)). However, whether LFP power in different frequency bands during neutral cueing is also intermediate between cued and uncued conditions is unknown.

While earlier neurophysiological studies focused mainly on responses of individual neurons, simultaneous recordings from multiple neurons have allowed a more comprehensive investigation of the effect of attention on the neuronal population. In particular, attention has been shown to reduce spike-count correlation between pairs of neurons (Cohen and Maunsell, 2009; Mitchell et al., 2009), which can affect the way the neural population encodes a stimulus (Averbeck et al., 2006; Cohen and Kohn, 2011). The spiking activity of a simultaneously recorded population of neurons has also been used to predict the animals’ focus of attention and behavior in a single trial (Cohen and Maunsell, 2010, 2011). Since handling high dimensional data is difficult, many studies have devised methods to obtain a one-dimensional metric from a high-dimensional dataset to predict behavior (Cohen and Maunsell, 2010; Cunningham and Yu, 2014; Yates et al., 2020). Using a simple method of projecting trials onto a one-dimensional axis constructed using the mean spiking activity of a neuronal population of two different attention conditions (Cohen and Maunsell, 2010; Mayo et al., 2015), Mayo and Maunsell, (2016) showed that the projections of neutral cueing condition were also intermediate between cued and uncued conditions. However, whether the projections remain similarly graded even when incorporating higher-order statistics like covariance across electrodes was not tested. Further, it is unclear how well different attentional states can be discriminated using LFP power at different frequency bands as compared to spiking activity, and how this discriminability varies with the number of microelectrodes.

To address these issues, we recorded spikes and LFPs from two macaque monkeys performing an attention task under different cueing conditions and compared how power in different frequency bands varied across attention conditions. We also compared LFP-LFP and spike-LFP coherence. Finally, we compared different weighting strategies to obtain one-dimensional projections of data using spikes or power in different frequency bands and compared the discriminability of attention conditions across neural measures as a function of population size.

## Materials and Methods

### Electrophysiological recordings

Data used in this study and the experimental procedures were detailed in a previous report (Mayo and Maunsell, 2016) that described the V4 spiking data in detail. All animal procedures were approved by the Institutional Animal Care and Use Committee of Harvard Medical School. Two adult male rhesus monkeys (*Macaca mulatta*) were surgically implanted with a titanium head post and scleral eye coil before training. After they learned the behavioral task, a 6 × 8 array of microelectrodes (Blackrock Microsystem, 48 electrodes in each) was implanted in visual area V4 in both cerebral hemispheres. The electrodes were 1 mm long and 400 μm apart with impedance typically between 0.2 and 1 MΩ at 1 KHz. Area V4 was identified using stereotaxic coordinates and by locating the lunate and superior temporal sulci during surgery. The azimuths and elevations of the centers of the receptive fields (RF) of neurons recorded from the two arrays were around (2°,-7°) and (−7°,-4°) for Monkey A, (6°,-7°) and (−6°,-3°) for Monkey W.

Signals from 96 channels were recorded using a 128-channel Cerebus Neural Signal Processor (Blackrock Microsystems). Signals were filtered between 0.3 Hz (Butterworth filter, 1^st^ order, analog) and 2.5 kHz (Butterworth filter, 4^th^ order, digital), sampled at 2 kHz and digitized at 16-bit resolution to obtain the LFP signals. Note that the low-pass LFP filter was inadvertently set to a value greater than half the sampling rate and hence could have led to some aliasing. However, because the power of the LFP signal falls rapidly with frequency (log PSDs shown in Figure 1C kept decreasing beyond 200 Hz to about −1.3 at ~700 Hz after which they flattened), the aliased power above 1000 Hz was negligible. Signals were separately filtered between 250 Hz (Butterworth filter, 4^th^ order, digital) and 7.5 kHz (Butterworth filter, 3^rd^ order, analog) and subjected to a threshold followed by spike sorting using spike sorter software (Plexon) to extract single and multi-units, which we also refer to as “neurons.”

**Figure 1:**
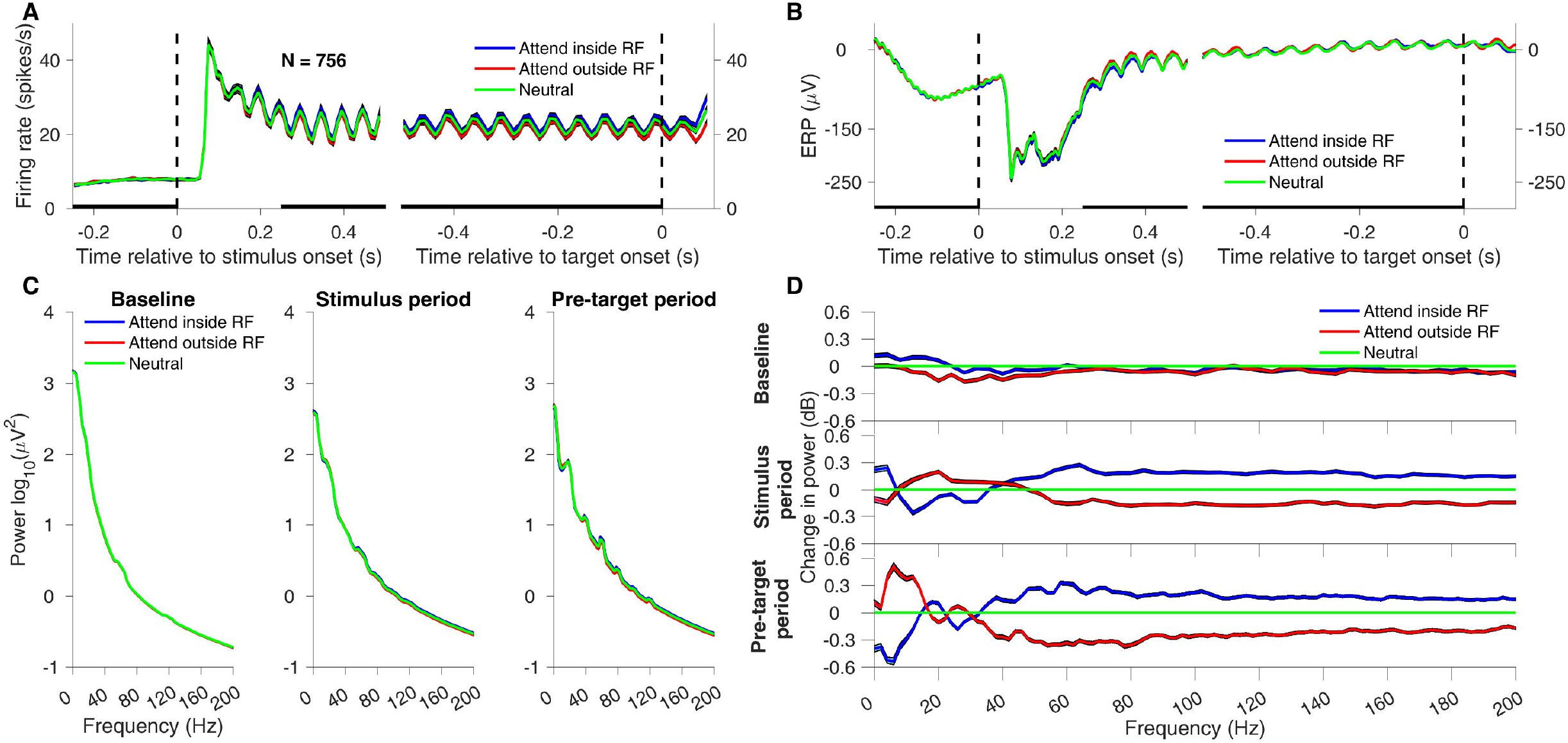
Comparison of firing rate, evoked potential and power spectral densities across different attention conditions. A) Mean PSTH of all 756 electrodes relative to stimulus onset (left plot) and target onset (right plot) for attend-in (blue), attend-out (red) and neutral (green) attention condition. The vertical dashed line indicates the stimulus onset time (left) and target onset time (right). The bars on the X-axis indicate the periods chosen for analysis. We only consider the valid and correct responses here (blue: Attend-in Valid Hit, red: Attend-out Valid Hit, green: Target-out Neutral Hit and Target-in Neutral Hit combined). Results for other attention conditions are shown in Figure 3. B) Mean Event Related Potential (ERP) of the same electrodes and periods as in (A) for the three attention conditions. C) Power Spectral Density of the baseline period (−0.25 to 0 s relative to stimulus onset), stimulus period (0.25 to 0.5 s relative to stimulus onset) and pre-target period (−0.5 to 0 s relative to target onset) for the three attention conditions. D) Change in power (in dB) of the three attention conditions, obtained by subtracting the green trace from the other traces shown in C. Green trace represents the change in power of the neutral condition with itself and hence is trivially zero.

### Behavioral task

Monkeys learned an orientation change detection task in which they held their gaze within a 1.8° square window centered on a fixation spot in the center of the screen throughout the trial. During fixation, two counterphasing Gabor stimuli appeared simultaneously and synchronously in the RFs of one of the units recorded from each hemisphere. At an unsignalled time, chosen from an exponential distribution (mean 3000 ms, range 500-5000 ms) and when the contrasts of the counterphasing Gabor stimuli were both at 0%, the orientation of one of the stimuli changed. Monkeys were rewarded for making a saccade to the location of the changed stimulus between 100 and 550 ms after the orientation change.

Monkeys were cued to attend to one (valid or invalid cue) or both (neutral cue) of the stimulus locations for a block of 50 trials using four instruction trials in which a white spot (0.45° radius) appeared for 50-100 ms indicating the likely location of the stimulus change in the upcoming cued block or simultaneously at both stimulus locations for the neutral block. In cued blocks, the change occurred at the cued location with 80% probability (valid cue) and 20% at the uncued side (invalid cue). In neutral blocks, the change occurred at each stimulus location with equal probability. Five percent of all trials were catch trials in which neither of the stimuli changed orientation and monkeys were rewarded for maintaining fixation throughout the trial. Catch trials and instruction trials were excluded from analysis.

Six randomly interleaved orientation change magnitudes were used in each session to characterize the effects of attention on behavioral performance. While all of these orientation changes were used during valid and neutral trials, only the second and third smallest orientation changes were used for invalid trials (see Mayo and Maunsell, 2016, for details). For these orientation changes, the mean behavioral performance was ~73% for validly cued trials, ~64% for neutral and ~48% for invalid trials. As in the previous report (Mayo and Maunsell, 2016), all analyses were restricted to only trials with these orientation changes for proper comparison of valid and neutral conditions with the invalid condition.

### Stimuli

Two odd symmetric, Gabor stimuli sinusoidally-counterphased at 10 Hz of maximum contrast were used during each recording session. Each had a size (σ range: 0.45°-1.6°) and spatial frequency (range: 0.3-2.5 cycles/deg) optimized for one of the units from each hemisphere. Stimuli were displayed on a CRT monitor (100 Hz frame rate, 1024 × 768 pixels, 8-bit DACs) over a uniform gray background. The monitor was calibrated to produce linear steps of luminance and positioned at 57 cm from the subject. Such counterphasing stimuli produce a salient neuronal response at twice the stimulus frequency (20 Hz; see Fig. 1A,B). These responses are referred to as steady-state visually evoked responses (SSVEPs). Eye position was sampled at 200 Hz using the scleral eye coil technique (Judge et al., 1980).

### Attention conditions

Attention conditions were defined in two ways. The first formulation was in reference to the RF of the neurons (Figures 3 and 4) and yielded the standard attend-in and attend-out conditions. The second formulation was used for population discriminability analysis that included neurons from both hemispheres (Figure 5) and was therefore based only on the cued hemifield. Each formulation consisted of 12 conditions, as described below.

First formulation: During the cued and uncued conditions, the animals’ attention was cued either inside or outside the RF. The cue could be either valid or invalid, and the animal could either detect (hit) or not detect (miss) the target. This yielded the following eight conditions: Attend-in Valid Hit, Attend-out Valid Hit, Attend-in Valid Miss, Attend-out Valid Miss, Attend-in Invalid Hit, Attend-out Invalid Hit, Attend-in Invalid Miss and Attend-out Invalid Miss. During the neutral cueing conditions, trials cannot be designated as attend-in and attend-out but can be separated based on where the final target appeared. This yielded the following four conditions: Target-in Neutral Hit, Target-out Neutral Hit, Target-in Neutral Miss, and Target-out Neutral Miss. Therefore, each attention condition was denoted by three words. The first word denoted the likely focus of attention/actual target location (two values: Attend-in/Attend-out or Target-in/Target-out); the second word denoted the validity of the cue (three values: Valid, Neutral or Invalid); and the third word indicated the behavioral outcome (two values: Hit or Miss).

Second formulation: This formulation was independent of RF location because electrodes from both hemispheres were used. Therefore, instead of using Attend-in or Attend-out as used in the previous formulation, we used Attend-left or Attend-right, where left/right indicated the attended visual hemifield.

### Session and electrode selection

Data were collected in 26 recording sessions (14 in Monkey A and 12 in Monkey W). One session was discarded because of an insufficient number of trials in one attention condition, yielding 25 sessions (13 and 12 in the two monkeys). For a fair comparison between spiking and LFP data, we only considered electrodes that had neuronal firing rates in the stimulus period that were >=5 spikes/s greater than during the baseline period. This yielded a total of 756 electrodes (average: 30.2 per session; min: 8, max: 46). Only sessions for which at least 15 trials were available for a particular attention condition were used for the analysis of that condition.

### Data analysis

Spike and LFP data were analyzed in three periods: (i) Baseline: 250 – 0 ms before stimulus onset, (ii) Stimulus: 250 – 500 ms after the stimulus onset, (iii) Pre-target: 500 – 0 ms before the stimulus changed to a different orientation (target). Peri-stimulus time histograms (PSTH) were obtained by binning spike counts in 10 ms windows. Power spectral density (PSD) was calculated using multi-taper method (Mitra and Pesaran, 1999) with three tapers using the Chronux toolbox in MATLAB (Bokil et al., 2010). Frequency resolution was 4 Hz in baseline and stimulus periods and 2 Hz in the pre-target period. Power in the specific bands was computed by adding the power at each frequency point in specified range for each electrode and then averaging across electrodes of all the sessions. The following bands were used: alpha (8-12 Hz), gamma (42-78 Hz) and high-gamma (122-198 Hz). The power at the harmonics of 20 Hz SSVEP was excluded from all the bands. The phase consistency between: 1) spikes and LFPs, and 2) LFPs and LFPs was computed across electrode pairs using pairwise phase consistency (PPC) which gives an unbiased estimate of the square of the phase consistency (Vinck et al., 2010, 2012).

### Population projection

The goal of the population projection analysis was to quantify how discriminable the population neural activity was when the monkey paid attention to the left versus the right stimulus location (i.e., the discriminability in the neural data between the Attend-left Valid Hit and Attend-right Valid Hit conditions). These two conditions were chosen because the animals were most likely to be paying attention to the cued locations when they detected a target on the validly cued side. Because the population data was high-dimensional (equal to the number of electrodes in both hemispheres), we wanted to find a one-dimensional projection (a weighted sum) of this high-dimensional data so that the projections were maximally separated for the two conditions. For this purpose, we used a linear discriminant analysis (LDA), which is a technique based on Fisher’s Linear Discriminant (Fisher, 1936) that maximizes the separation between two classes in a lower-dimensional space. This method works well when the size of the test data (number of trials in the Attend-in and Attend-out Valid Hit conditions) is comparable to the dimensionality (number of electrodes), or if assumptions of common covariance matrix across classes are not met (Li et al., 2006). More complex classifiers like neural networks were not considered here because they typically require a larger training dataset set to prevent overfitting, and they are less interpretable.

We used three projection methods of increasing complexity, progressively building to the LDA because this allowed us to study the effect of accounting for mean differences across classes, variances and covariances separately (see below for details). Further, we wanted to also compare with a technique called “Attention Axis” analysis (figure 5; Cohen and Maunsell, 2010; Mayo et al., 2015), which is a variant of the first method. All analyses were performed both with and without 5-fold cross-validation to better illustrate the effect of cross-validation on our results.

Consider two data matrices, D_L_ and D_R_, corresponding to the neural data for the Attend-left Valid Hit and Attend-right Valid Hit conditions. The sizes of these matrices are NxT_L_ and NxT_R_, where N is the total number of electrodes from both hemispheres, and T_L_ and T_R_ are the number of trials for the left and right conditions. Our aim is to find an Nx1 weight vector W, which is used to obtain the projections P = W^T^D. Although W is obtained using D_L_ and D_R_ only, it can be used to project the data (D) of any attention condition. The three methods are as follows:

1. Mean-Difference: Here *W* = (*μ_L_* — *μ_R_*), where *μ_L_* and *μ_R_* represent the mean of the data matrices D_L_ and D_R_ across trials respectively and therefore are Nx1 matrices. Here, we assign a weight to each electrode that depends on how different its responses are for the two attention conditions. If the responses are equal for the two attention conditions, the weight for that electrode is zero. The first step of the “Attention Axis” projection is similar to this step. This method does not account for the variance across trials for individual electrodes or the covariances across electrodes.
2. Uncorrelated LDA: Here, we divide each weight obtained in the previous case by the average weighted variance across trials for the two conditions. For the i^th^ electrode, the weight is given by:

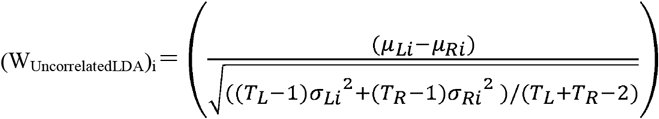 Where *μ_Li/Ri_* is the mean activity for the i^th^ electrode as before, while *σ_Li/Ri_*^2^ is the variance in neural activity for left/right condition for the i^th^ electrode. In vector form, W_UncorrelatedLDA_ = *∑*^−1^ (*μ_L_* – *μ_R_*), where *∑* is an NxN diagonal matrix containing the pooled variances of the electrodes along the diagonals.
3. LDA: To also account for co-variations in neural data across electrode pairs, we use the full covariance matrix instead of a diagonal matrix:

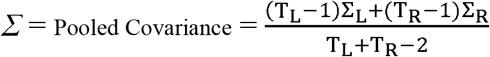 Where ∑_L/R_ is the NxN covariance matrix of D_L/R_ and W_LDA_ = *∑*^−1^ (*μ_L_* – *μ_R_*) as before.

### Simulations

To study the effects of cross-validation, regularization, and sample size in LDA analysis, we generated two datasets of 50000 samples, each drawn from two 30-dimensional multivariate normal distributions with different mean vectors and the same covariance matrix such that the true d-prime (ratio of the difference of the means and the pooled variance) of the projection of data onto the optimal LDA axis was between 0.5 and 1.5. We then randomly drew samples of different sizes from these datasets and computed the d-prime of projections onto the LDA axis estimated with or without 5-fold cross-validation for each sample size. Finally, we normalized the estimated d-prime by taking the ratio with the true d-prime. This process was repeated 50 times to compute the standard deviations.

These simulations highlight potential issues with neurophysiological datasets in which the number of dimensions (electrodes, ~30 on average in our case) is not much smaller than the number of data points. In such cases, not cross validating leads to severe overfitting, while cross-validation leads to poor performance because it is difficult to estimate a large covariance matrix and the means of each class with few data. For example, we noticed in our simulations that when the number of data points per class (trials per attention condition) was small (say, ~100 trials), the estimated d-primes were ~1.51 ± 0.03 (mean ± SEM) when not cross-validated, but only ~0.6 ± 0.03 when cross-validated. Only when the number of trials was several folds (>10 times) larger than the number of dimensions was the estimated d-prime close to the true d-prime.

If the number of trials is less than the dimensionality, the covariance matrix is no longer invertible and needs to be “regularized” (Friedman, 1989; Guo et al., 2007). The *fitcdiscr* function in MATLAB that was used to perform the LDA analysis applies some regularization by default, but it can be further modified by using the *cvshrink* function in MATLAB. The modification of the covariance matrix depends on the parameter *gamma*, which ranges from 0 to 1, which shrinks the covariance matrix towards a diagonal matrix. In our simulations, we used 30 different gamma values and chose the one that minimized the classification error. We also used another type of regularization where an additional parameter *delta* was varied along with *gamma* that sets a threshold for choosing the useful predictors (dimensions). However, varying *delta* did not give any advantage over varying just *gamma* (data not shown), and varying gamma also did not improve the performance over the default regularization performed by *fitcdiscr* (Supplementary Figure 2). For real data, therefore, we did not do any specific regularization.

## Results

Neural recordings were made using two 6 × 8 (48 channels) arrays implanted bilaterally in visual area V4 of both hemispheres in two rhesus monkeys (*Macaca mulatta*). Figure 1A shows the average Peri-Stimulus Time Histogram (PSTH) of 756 neurons from 25 recording sessions in two monkeys, aligned to either the stimulus onset (left vertical dashed line) or target onset (right vertical dashed line). Here, attend-in and attend-out conditions are for validly cued stimulus changes that the monkeys correctly detected (Attend-in Valid Hit and Attend-out Valid Hit; see Methods for details), while the neutral condition includes all neutrally-cued trials in which the monkeys detected the target, irrespective of target location (Target-in Neutral Hit + Target-out Neutral Hit; see Methods for details). There was a transient response after stimulus onset, followed by a 20 Hz SSVEP in the PSTH due to the 10 Hz counterphase stimulus. Throughout each trial attention state had a small effect on spiking, with activity during the neutral condition (green trace) below attend-in (blue) and above attend-out (red) conditions, as reported previously (Mayo and Maunsell, 2016).

Figure 1B shows the event related potential (ERP), obtained by averaging the raw LFP traces across trials, of the same three attention conditions. Like the PSTH, the ERP had a stimulus onset-related transient and a prominent 20 Hz SSVEP, with the neutral condition (green) between the attend-in and attend-out conditions. Figure 1C shows the power spectral density (PSD) of the attention conditions for baseline, stimulus and pre-target periods (indicated as thick horizontal black lines in Figure 1A). In all conditions, the PSDs showed a typical “1/f” power-law decay. The PSDs for the baseline condition were largely overlapping. In the stimulus-onset and pre-target conditions, there was a small separation in the three traces, with the neutral (green) trace almost always in between the attend-in and attend-out conditions. These PSDs also showed prominent SSVEP bumps at 20 Hz and its harmonics. These SSVEP peaks were best observed in the pre-target period because of better frequency resolution (2 Hz, compared to 4 Hz for the other two periods). These plots show that the effect of attention on PSDs is weak at an absolute scale.

Figure 1D shows the change in power in the attend-in and attend-out condition relative to the neutral attention condition for the three periods, obtained by subtracting the green trace from the other traces. Because PSDs are shown on a log scale in Fig. 1C, subtraction on a log scale is simply the log of the ratio of power change across conditions (after subtraction, the log ratio is multiplied by 10 to get units of decibels). Previous studies have shown that attention suppresses power at low frequencies and increases power at higher frequencies (Fries et al., 2001; 2008). Consistent with this, we observed that the power in lower frequencies was suppressed for attend-in condition relative to attend-out and the effect reversed at higher frequencies (>30 Hz), with the neutral condition almost always between the two. This effect was particularly pronounced during the pre-target period (Fig. 1D, bottom row), where the sampled brain activity presumably reflected the animals’ state of attention most accurately before the impending, successfully detected, target change. The increase in power at high frequencies was broadband and extended up to 200 Hz and beyond, with a slight peak in the gamma range (30-80 Hz) in the pre-target condition. An increase of 0.5 dB corresponds to a change in 10^0.05^ = 1.12, or 12% increase in power, comparable to the size observed in firing rates in previous spatial attention studies (Boudreau et al., 2006; Mayo and Maunsell, 2016; Snyder et al., 2018). These changes were highly reliable across electrodes and sessions, yielding very small standard error of mean (shaded region around the traces). As shown below, this allowed better discrimination between different attention conditions using LFP power compared to spiking activity.

Previous studies have suggested that phase synchronization between LFPs from two electrodes as well as between spikes and LFPs increase in the gamma range with spatial attention in area V4 (Fries et al., 2008; Fries, 2015). We therefore measured phase consistency across electrode pairs using pairwise phase consistency (PPC), an unbiased estimator of the square of the phase coherence. Figure 2A shows the field-field (LFP-LFP) PPC across all pairs of selected electrodes within the hemisphere. As before, the neutral condition was between the attend-in and attend-out at almost all frequencies. Peaks were observed at 20 Hz and its harmonics because the counterphasing stimulus forced a phase consistency across electrodes. Figure 2B shows the change in PPC from the neutral condition, obtained by subtracting the green trace from each of the traces in Figure 2A. In contrast to the change in power (Figure 1D), the change in PPC was prominent mainly in the gamma range (30-80 Hz), with negligible differences at higher frequencies (>120 Hz). This could be because phase synchronization measures are less sensitive at high frequencies because even tiny absolute shifts in the time domain correspond to large shifts in phase, leading to poorer phase consistency in the presence of small time shifts (Ray, 2015). The spike-field (spike-LFP) PPC (Figure 2C) showed strong peaks at 20 Hz and harmonics, indicating that spikes were tightly locked to the SSVEP. The change in PPC relative to the neutral condition (Figure 2D) showed similar trends to that in field-field PPC, but the magnitude of the effect was far smaller. There was also a reduction in phase coherence with attention at very low frequencies (<5 Hz) in both field-field and spike-field coherence, possibly due to the reduction in power at those frequencies with attention.

**Figure 2:**
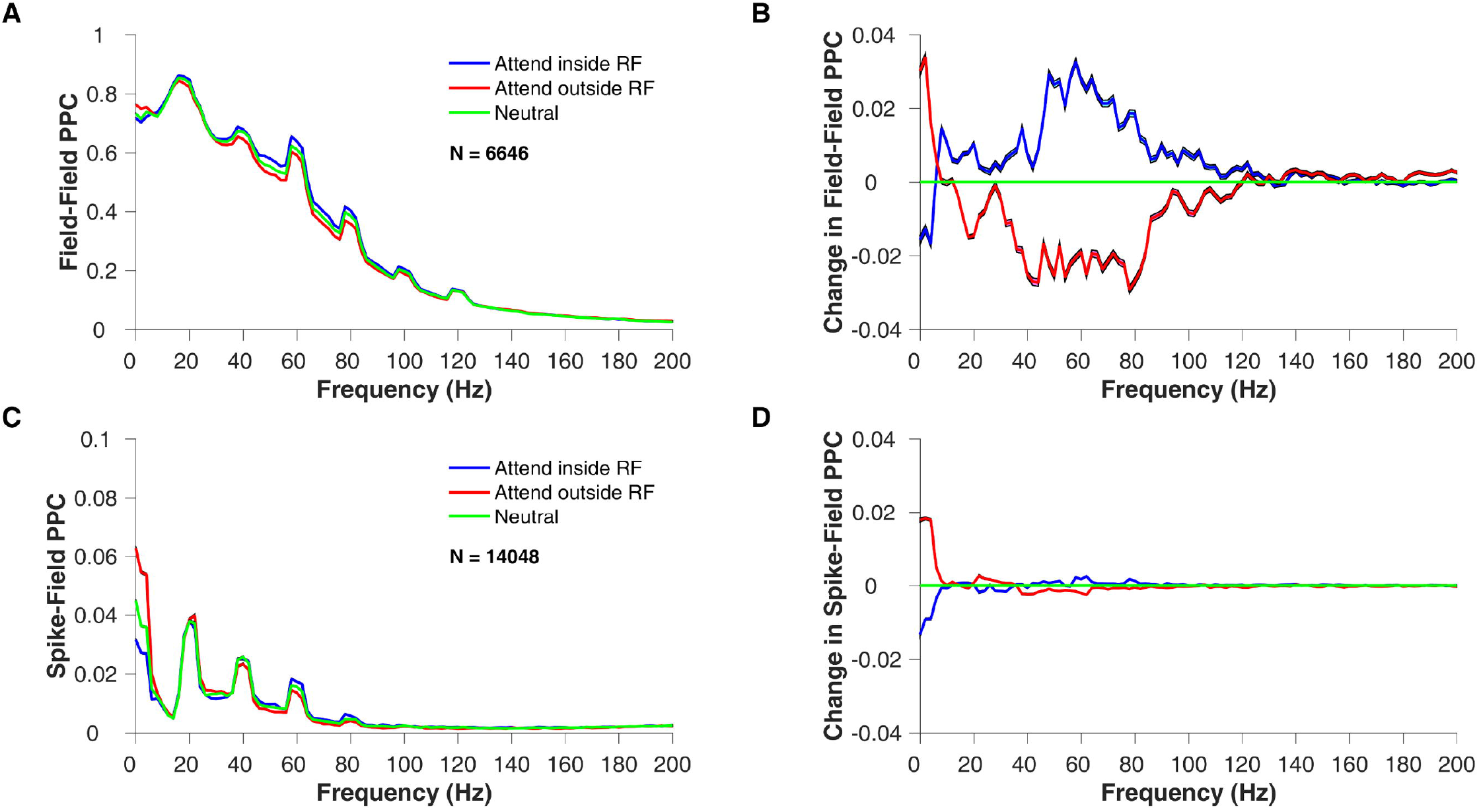
Comparison of intra-hemispheric Field-field (LFP-LFP) and Spike-field (Spike-LFP) phase relationship across attend-in, attend-out and neutral attention conditions. **(A)** Mean field-field pairwise phase consistency (PPC), averaged across 6646 electrode pairs of attend-in (blue trace), attend-out (red) and neutral condition (green). **(B)** Mean change in field-field PPC of the three attention conditions, relative to the neutral condition. As before, the neutral condition is trivially zero. **(C)** Mean Spike-field pairwise phase consistency (PPC) averaged across 14048 electrode pairs (including spike and LFP from the same electrode) for the three attention conditions. **(D)** Mean change in Spike-field PPC for the three attention conditions, relative to the neutral condition.

We also studied the phase relationship between the LFP signals across hemispheres and observed peaks at 20 Hz SSVEP and its harmonics, as in the within-hemisphere case, in both field-field and spike-field PPC, although the magnitude was reduced (Supplementary figure 1A and C). In this case the electrodes of each across-hemisphere pair have RFs in opposite visual hemifields, there were no “attend-in” and “attend-out” conditions; so we instead took the average of attend-left and attend-right conditions and compared them with the neutral condition. The difference between attend-left/attend-right and neutral conditions showed inconsistent trends (Supplementary figure 1 B and D) with small peaks at 20Hz SSVEP and harmonics.

Our main goal was to compare how well the animals’ behavior could be predicted using spiking activity versus LFP power. To see the consistency of effects, we performed this comparison over 12 different attention conditions (see Materials and Methods for details). We first compared how firing rates and LFP power at different frequencies varied across different attention conditions. Rather than using LFP power at individual frequency bins, we computed LFP power in three popular frequency bands – alpha (8-12 Hz), gamma (42-78 Hz) and high-gamma (122-198 Hz). Averaging of power within a frequency range is needed because single-trial estimates of power at individual frequency bins are noisy (Jarvis and Mitra, 2001; Srinath and Ray, 2014; Chandran KS et al., 2018). However, because the noise in spectral estimation is largely independent across frequencies, averaging power within a large frequency range provides reliable estimates of the actual power on single trials. The lower limit of high-gamma range was set at a higher value than usual (122 Hz instead of 82 Hz) because we wanted a frequency range in which field-field coherence was close to zero and therefore unlikely to simply reflect the dynamics of the traditional gamma band. However, choosing the high-gamma range between 82-198 Hz yielded similar results.

Figure 3A shows the average firing rates (top row), as well as raw power in the high-gamma (second row), gamma (third row), and alpha ranges (bottom row) for 12 different attention conditions. The conditions are grouped based on cue validity (left column: valid, middle column: neutral and right column: invalid), and within each group, conditions are arranged such that the firing rate was expected to increase going from left to right, as discussed below.

**Figure 3:**
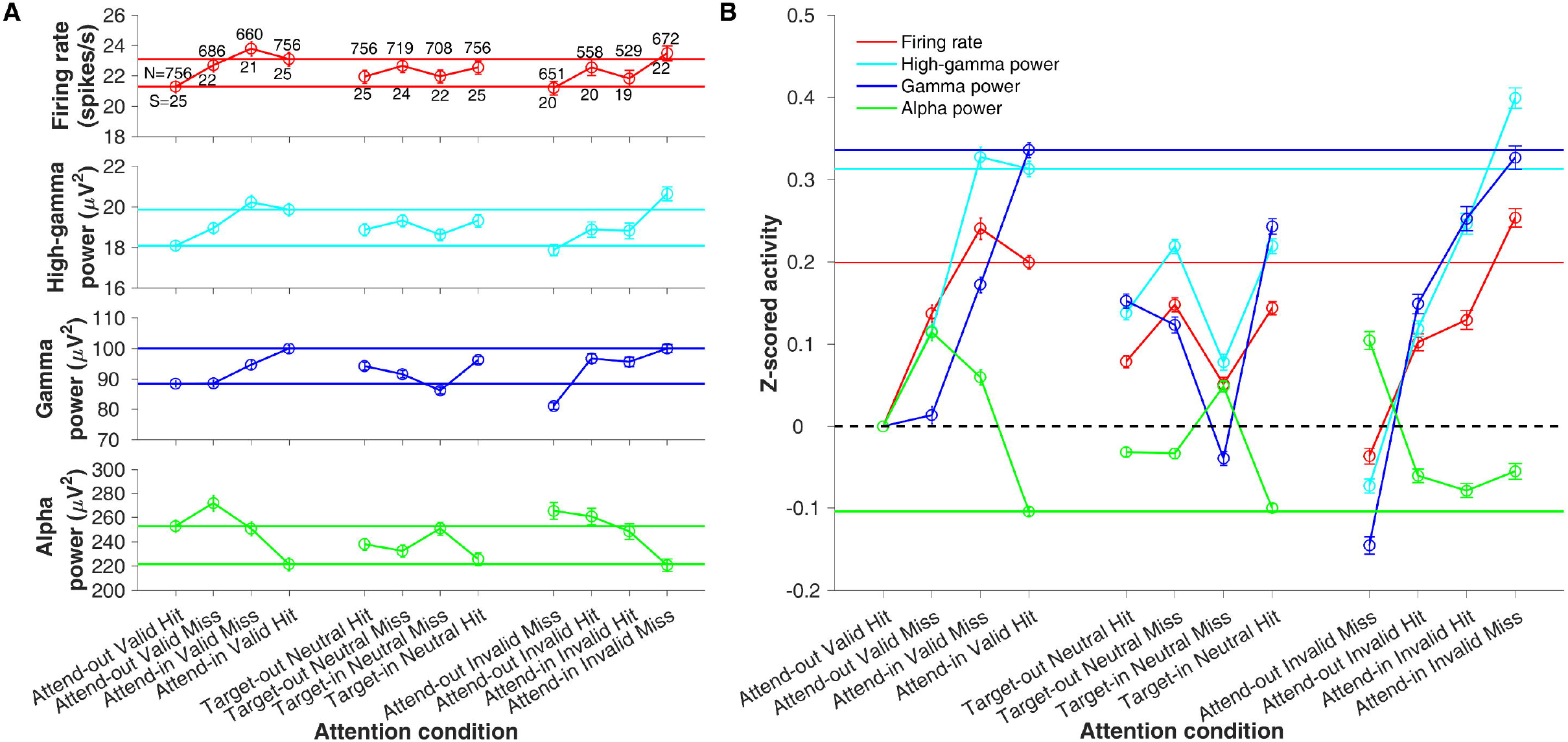
Comparison of firing rate, high-gamma power, gamma power, and alpha power across 12 attention conditions. A) First row: mean firing rate during the pre-target period for the Hit and Miss trials where attention was validly cued out and into the RF (left column); for Hit and Miss trials of neutral condition where the target appeared out and in the RF (middle column); and for Hit and Miss trials where attention was invalidly cued out and into the RF (right column). The numbers above the error bars indicate the number of electrodes averaged across for the respective conditions, while the numbers below the error bars indicate the number of sessions. We only considered sessions that had more than 15 trials for an attention condition. Second, third and fourth rows: mean high-gamma, gamma and alpha power, respectively. The same number of electrodes and sessions were used for these measures as well. Error bars indicate the s.e.m. The validly cued attend-out and attend-in conditions are indicated by horizontal lines for better comparison with the other conditions. B) Same data as A, but after z-scoring the data of each electrode and attention condition by subtracting the mean and dividing by the standard deviation (computed across trials) of the attend-out valid hit condition. The colored horizontal lines indicate the validly cued attend-in values for each neural measure. The black horizontal dashed line indicates the validly cued attend-out condition, which is trivially zero.

For the valid conditions (Figure 3A; left column), firing rates were lower for Attend-out Valid Hit (first data point) than Attend-in Valid Hit (fourth data point), as shown previously (Figure 1). Similar trends were observed in high-gamma and gamma power, while alpha power showed the opposite result. For the same attentional conditions, the monkeys were expected to miss the target if their attention was deployed on the “wrong” side, and therefore the corresponding miss conditions (second and third data points) were expected to lie between the two hit conditions. With a few exceptions, this was the case.

For the neutral case (Figure 3A; middle column), the attention conditions were divided based on the trial outcome and target location rather than cue location since monkeys were cued to attend to both locations. Like the changes in firing rate, high-gamma power and gamma power were lower when monkeys detected the target that appeared outside the RF than when it appeared inside the RF, while alpha power showed the opposite trend. As before, barring a few exceptions, missed conditions lay between the hit conditions for all the neural measures. Importantly, almost all the neutral attention conditions lay between validly cued attend-out and attend-in conditions (marked by solid horizontal lines for clarity).

In the invalid case (Figure 3A; right column), the target always appeared at the uncued location. Firing rate, high-gamma, and gamma power were lowest when attention was directed outside the RF and the monkey missed the target, as expected (first data point). Likewise, these measures were highest when the monkey attended inside the RF but missed an invalidly cued stimulus change outside the RF, likely because attention remained focused on the unchanged stimulus within the RF (fourth data point). These two miss conditions were comparable to the Attend-out Valid Hit and Attend-in Valid Hit conditions respectively regarding the focus of attention. As before, alpha power showed the opposite effect. The hit trials, for which the monkeys were likely attending to the uncued side where the target actually appeared, lay between the missed trials for the invalid condition for all the neural measures.

These results show that both firing rate and LFP power in different frequency bands vary systematically with the attentional state, and therefore could potentially be used to predict the animals’ behavior. To directly compare the discriminability of different neural measures, we z-scored all measures separately for each electrode and attention condition, by subtracting the mean and dividing by the standard deviation across trials of the Attend-out Valid Hit condition. The z-scored values therefore indicated how far a particular attention condition was from the Attend-out Valid Hit condition, in units of the standard deviation of Attend-out Valid Hit condition across trials. This condition was chosen for simplicity: the discriminability remained similar if we normalized all measures relative to Attend-in Valid Hit condition or used the average standard deviation of the Attend-out and Attend-in Valid Hit conditions.

Figure 3B shows the mean z-scored values (averaged across electrodes) for different attention conditions for all four neural measures, as indicated by line colors. The difference in z-scored activity between Attend-out Valid Hit (which was zero by definition) and Attend-in Valid Hit was greatest for gamma power (z = 0.34 ± 0.0091; mean ±1 s.e.m), followed by high-gamma (0.31 ± 0.0094), firing rate (0.2 ± 0.0083) and finally alpha (−0.1 ± 0.0051). The difference in z-scores between gamma and high-gamma was not significant (p = 0.24; Wilcoxon rank sum test), but the difference was highly significant between high-gamma and firing rate (p = 2.3×10^−21^; Wilcoxon rank sum test) as well as gamma and firing rate (p = 3.84×10^−27^; Wilcoxon rank sum test). On the other hand, magnitudes of the z-scored values for alpha power were significantly less than firing rates (p = 2.68 ×10^−10^; Wilcoxon rank sum test). Therefore, if a single electrode is available for analysis, gamma/high-gamma power allows superior discrimination of attend-in and attend-out conditions compared to firing rates.

Previous studies have shown that when firing rates are recorded from multiple electrodes simultaneously, a suitable linear combination of these signals can substantially improve the discriminability across attentional conditions, and this discriminability further depended on the correlation between the total number of spikes recorded from two electrodes (“spike-count” correlation; Cohen and Maunsell, 2009). Analogous correlations between LFP power across electrode pairs have not been compared, to our knowledge, with spike-count correlations (but see (Snyder et al., 2015) for related work). We found that the absolute Pearson correlation between power recorded from two electrodes within the same hemisphere was highest for alpha power (Attend-out Valid Hit condition: 0.84 ± 0.0015), progressively decreasing for gamma and high-gamma power (0.69 ± 0.0018 and 0.46 ± 0.0019), and lowest for spike-count correlations (0.22 ± 0.002). However, the relative change in correlation from the Attend-out Valid Hit condition, as shown in Figure 4A, showed similar trends across the four neural measures. For valid cues (left column), spike-count correlations decreased from attend-in to attend-out conditions, consistent with previous studies (Cohen and Maunsell, 2009; Mitchell et al., 2009). Interestingly, we found that correlation of high-gamma, gamma, and alpha power were also reduced in Attend-in Valid Hit condition compared to Attend-out Valid Hit. As before, the “missed” cases showed intermediate correlations between the validly cued hit conditions in most cases.

**Figure 4:**
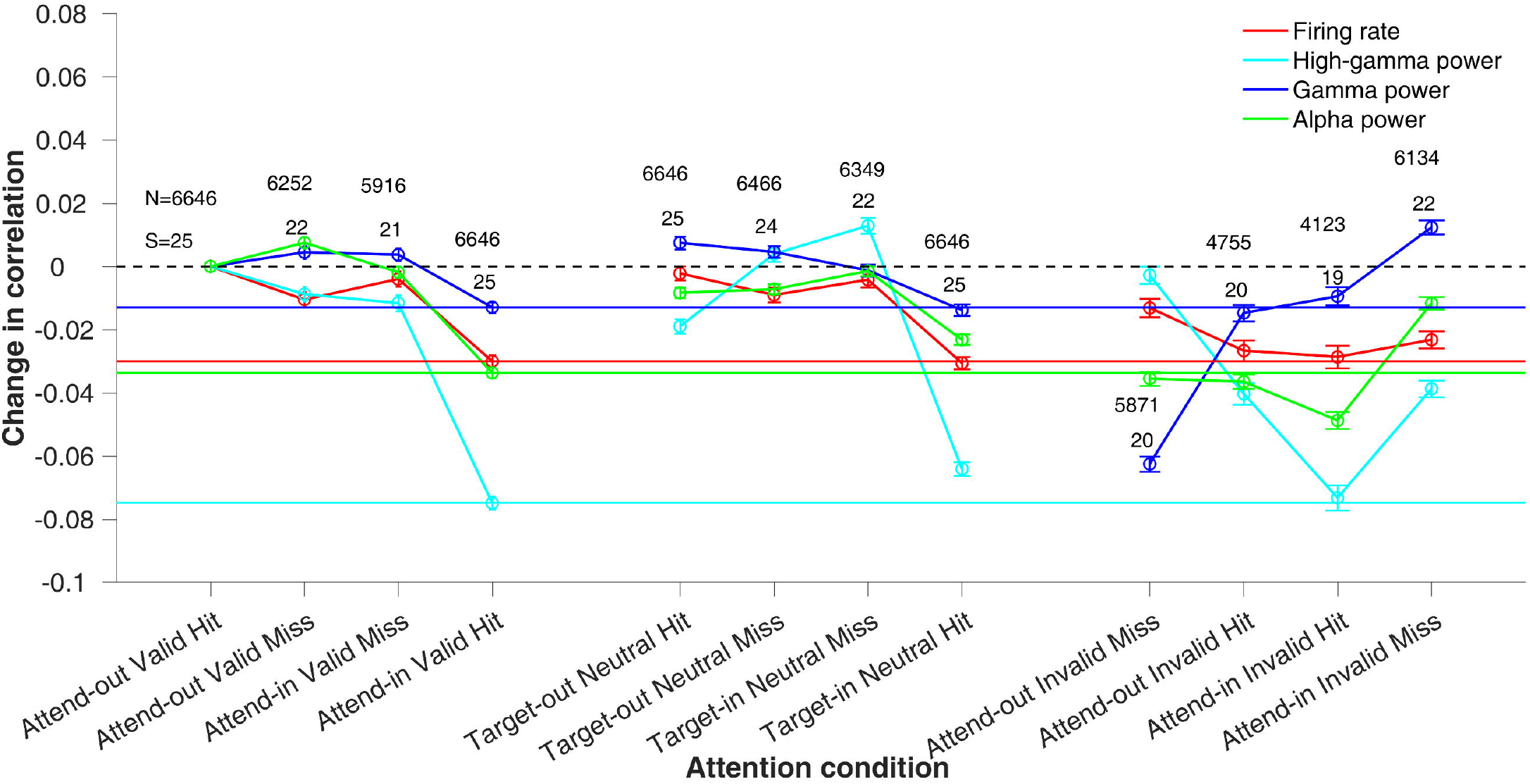
Intra-hemispheric Pearson correlation of firing rate, high-gamma, gamma and alpha power. Mean change in correlations between intra-hemispheric electrode pairs for the four neural measures relative to attend-out valid hit condition (same format as figure 3B). Numbers above the error bars of the blue line indicate the total number of electrode pairs (N) and sessions (S) considered for each condition. As in figure 3B, the colored horizontal lines indicate the validly cued attend-in values for each neural measure. The black horizontal dashed line indicates the validly cued attend-out condition, which is trivially zero.

For the neutral conditions (middle column), Mayo and Maunsell (2016) showed that spike-count correlations were intermediate between attend-in and attend-out. However, in their data, the correlations were computed for all correct trials, irrespective of the target position (in other words, Target-out Neutral Hit and Target-in Neutral Hit conditions were pooled, as done in Figures 1–2 here as well). However, when we computed the correlation separately for these two Hit conditions, we found that spike-count correlations for Target-out Neutral Hit (0.218 ± 0.002) were comparable to Attend-out Valid Hit (0.22 ± 0.002), although the difference was significant (p = 0.007; Wilcoxon rank sum test). Similarly, Target-in Neutral Hit (0.189 ± 0.002) were similar to Attend-in Valid Hit (0.19 ± 0.002, p=0.64; Wilcoxon rank sum test). The Miss conditions were again intermediate between the two Hit conditions in most cases. Finally, the invalid conditions (right column) showed predictable trends for firing rates, but variable results for the LFP power, especially correlations in gamma power that increased between Attend-out Invalid Miss versus Attend-in Invalid Miss. Potential reasons for this surprising effect are addressed below in the Discussion. Inter-hemispheric correlations were relatively weak (absolute correlation values for the valid hit condition were 0.273 ± 0.0018, 0.167 ± 0.0014, 0.104 ± 0.0015, 0.044 ± 0.0012 for alpha power, gamma power, high-gamma power and firing rate respectively) and difficult to interpret (data not shown).

Although the analysis shown in Figure 3 suggests that high-frequency LFP power is more discriminable than firing rates, this analysis only reflects the average discriminability computed from individual electrodes. Previous studies have shown that while single-channel LFPs outperform single-channel spikes, decoding based on multi-channel spikes is often superior to multi-channel LFPs ((Hwang and Andersen, 2013); see Discussion for details). This is because spiking activity is more dissimilar across electrodes as compared to LFP power (spikes have lower absolute correlations, as seen in our dataset as well) and consequently, discriminability between different attentional states is more variable across electrodes for firing rates than LFP power. Therefore, a weighting strategy that gave more weight to the more discriminable electrodes could improve the overall discriminability of the pooled signal (Jazayeri and Movshon, 2006; Graf et al., 2011), and improve the overall population discriminability of firing rates compared to LFP power. The optimum weights for a linear decoder can be obtained using Linear Discriminant Analysis (LDA). We used three pooling strategies (see Methods) to illustrate three factors (mean, variance and covariance) that are used in LDA analysis. We show these results first without any cross-validation since the effect size is larger in that case (Figure 5A). Results after 5-fold cross-validation are shown in Figure 5B.

**Figure 5:**
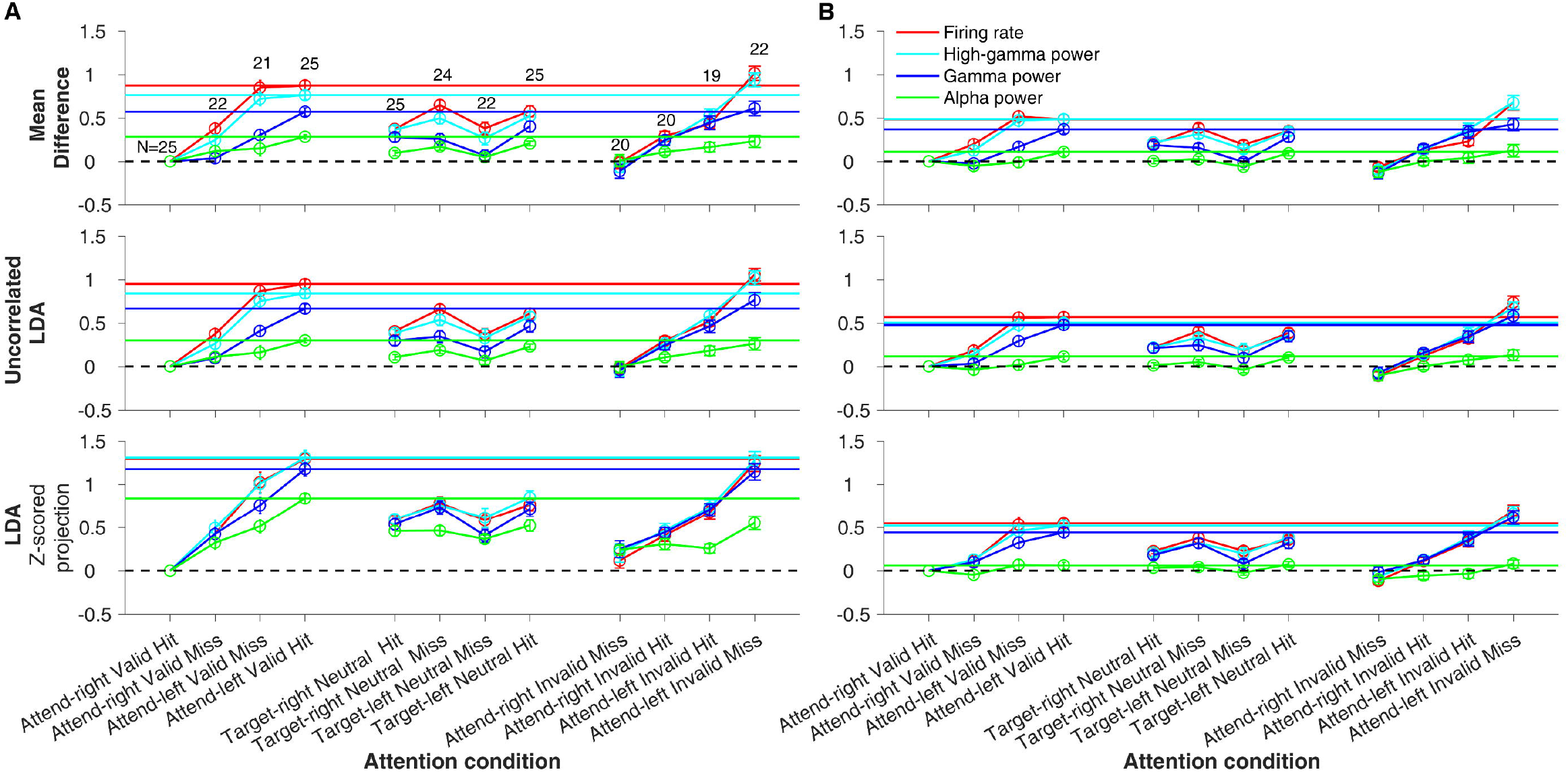
Comparison of projections onto mean difference, uncorrelated LDA and LDA weight vectors for different attention conditions. (A) Mean z-scored projections of firing rate (red trace), high-gamma power (cyan), gamma power (blue), alpha power (green) of trials of each attention conditions using non-cross validated weight vectors computed using mean difference (top row), uncorrelated LDA (middle row) and LDA (bottom row) methods (see text for details). The horizontal lines indicate the mean z-scored projections of the attend-left valid condition of the four neural measures (represented by their respective colors). The horizontal black dashed line indicates the z-scored projection of attend-right valid hit condition, which is trivially zero since z-scoring is done relative to that condition (see methods). The numbers on top of the data points in the top row indicate the number of sessions as before. Error bars indicate the s.e.m. (B) Same as that of (A) but with 5-fold cross-validation.

In the first pooling strategy (Figure 5A), we assigned a weight to each electrode that was equal to the difference in neural activity between the Attend-left Valid Hit and Attend-right Valid Hit conditions. The weighted average of all electrodes was then taken to obtain the projections for individual trials. The weights were defined such that attention to the left side led to larger projections than attention to the right side. The “Attention Axis” approach, which involves taking an inner product of the data vector with a vector created by connecting the mean of the data points for the Attend-left Valid Hit and Attend-right Valid Hit conditions, essentially performs the same operation (Cohen and Maunsell, 2010). To facilitate unbiased comparisons in discriminability across our four neural measures, we normalized all the projections by subtracting the mean value of the projections for the Attend-right Valid Hit condition and dividing by the standard deviation of the same condition (essentially z-scoring, as done in Figure 3B as well). Therefore, the mean projections for the Attend-right Valid Hit case were zero by definition, while these values were positive for Attend-left Valid Hit conditions, with larger values corresponding to better population discriminability.

Figure 5A (first row) shows the projections for all 12 attention conditions. Note that the computation of the weights was done using only 2 of these 12 conditions, the first and fourth data points in the left column, and subsequent z-scoring was done relative to the first data point. For the Attend-left Valid Hit condition, firing rates showed the largest value (0.87±0.06), followed by high-gamma (0.76±0.05; but the difference between the two was not statistically significant, p = 0.16, Wilcoxon rank sum test), gamma (0.57±0.05, significantly smaller than firing rates, p = 1.67×10^−4^ and high-gamma, p = 0.003, Wilcoxon rank sum test), and alpha power (0.28±0.03, significantly smaller than all three measures; firing rates: p = 3.21×10^−8^, high-gamma: p = 1.06×10^−7^, gamma: p = 8.1×10^−6^, Wilcoxon rank sum test). The Valid Miss conditions (second and third data points in the left column) were intermediate between the two Valid Hit conditions in all cases, as were the projections for all the Neutral conditions (middle column). Invalid conditions (right column) showed similar results as the Valid (with the Miss and Hit labels flipped, as in Figs. 3B and Figure 4).

The results were similar when we scaled individual weights by their variances (second row). However, when we also accounted for the covariances in the computation of weights (as done in LDA), we observed a large increase in the projection values for gamma and alpha power for which the absolute correlation was high (third row). Surprisingly, the projections of the Attend-left Valid Hit condition using gamma power (1.18±0.08) were now statistically indistinguishable to high-gamma (1.3±0.08) and firing rates (1.29±0.07) (p = 0.45, Kruskal-Wallis test for comparison of the three measures). This suggests that using high-frequency LFP is equally useful in discriminating between attention conditions as firing rates, even after using population data from several microelectrodes. Comparable changes in discriminability across neural measures were also observed for the Invalid conditions (right column), while the Neutral conditions had intermediate projections (middle column).

The results in Figure 5A were obtained without any cross-validation, which tends to inflate the discriminability (Subramanian and Simon, 2013) because the training and testing data are the same, which leads to overfitting. Figure 5B shows the same results using 5-fold cross-validation in which data were divided into five equal parts, out of which the weight vector was obtained using four parts (training data) and the projections were computed for the trials in the remaining part (testing data). This was repeated over by considering different testing set exhaustively among those five parts and then averaging across them. The results were similar to Fig 5A, although now there was not much difference between the three methods. For example, the mean projection values for Attend-left Valid Hit condition for firing rates were 0.48 ± 0.06, 0.57 ± 0.06 and 0.55 ± 0.05 (p = 0.45, Kruskal-Wallis test) for mean difference, uncorrelated LDA, and LDA, respectively. We show using simulations that this reduction in the LDA projection values was because the number of trials was not substantially larger than the number of dimensions (electrodes), especially after cross-validation for which the number of training trials was 80% of the total (Supplementary Figure 2). Despite this problem that led to lower projection values, the overall trends remained similar, with firing rate projections for the Attend-left Valid Hit (0.55 ± 0.05) not significantly different from high-gamma (0.52 ± 0.06) and gamma (0.44 ± 0.06) (p=0.5; Kruskal-Wallis test for comparison of the three measures). Perhaps because alpha power showed the weakest effects without cross-validation, it showed severely limited discriminability after cross-validation. As before, the invalid condition (right column) yielded similar results as valid (left column), while neutral condition (middle column) yielded intermediate values.

Finally, we studied how discriminability varied with the number of available electrodes. Figure 6 shows the z-score values of the Attend-left valid hit condition with cross-validation as a function of the number of electrodes, obtained by first selecting a subset of electrodes without replacement and performing the full LDA analysis (third pooling strategy) as before on this subset. When the number of electrodes was small, high-gamma and gamma power performed significantly better than the firing rate and alpha power (similar trends as in Figure 3B). As the number of electrodes increased to ~12, the discriminability of firing rate became comparable to gamma and high-gamma power. The discriminability reached an asymptote after ~15 electrodes, but this could be due to poorer performance of LDA with increasing dimensionality (Supplementary Figure 2). Discriminability also increased for alpha power for up to a few electrodes before reaching an asymptote but remained much lower than the other three measures. Overall, these results show that high-frequency LFP power is more useful than spiking activity when few electrodes (roughly 10-12 electrodes) are available and equally useful as firing rate with higher number of electrodes. Results were qualitatively similar without cross-validation, although the absolute values were larger (Supplementary Figure 3).

**Figure 6:**
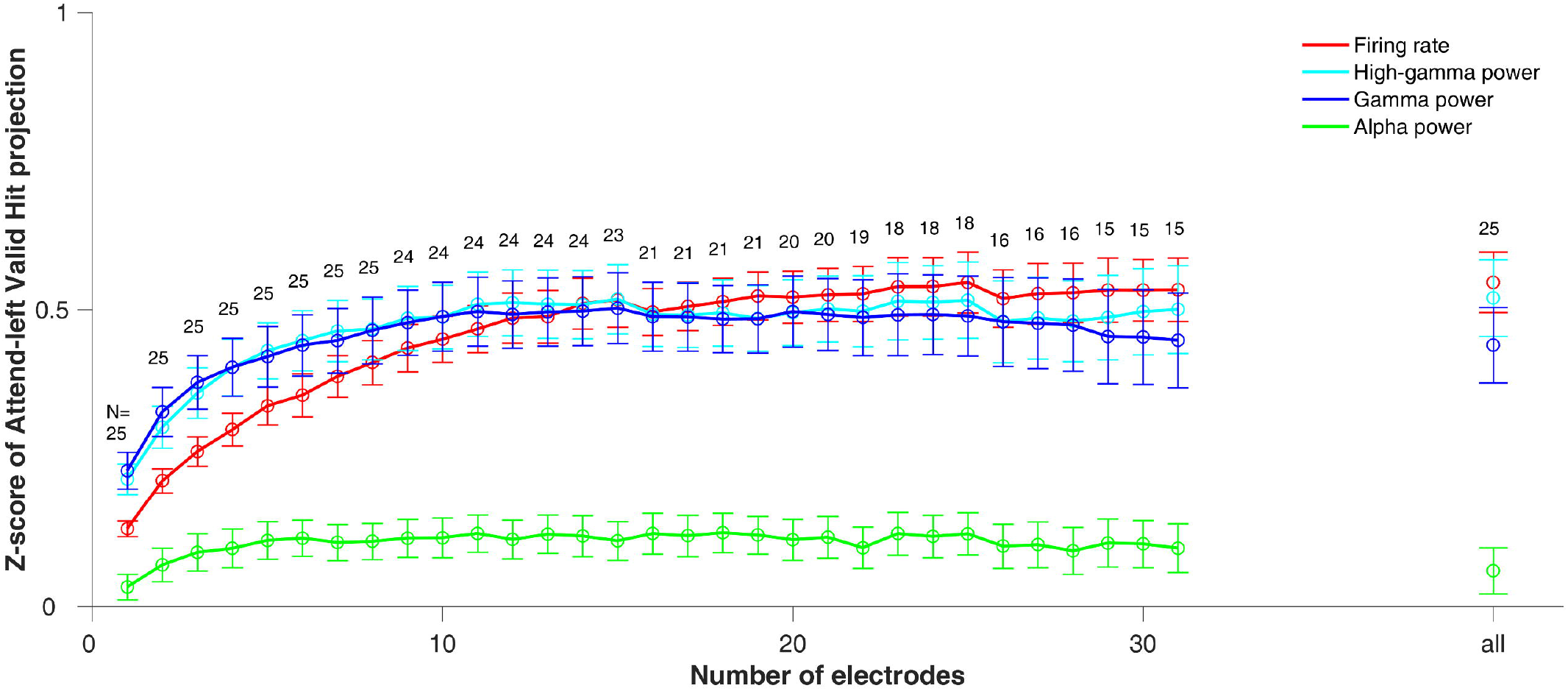
Discriminability as a function of the number of electrodes. Mean z-score of 5-fold cross-validated LDA projection of firing rate (red), high-gamma (cyan), gamma (blue) and alpha power (green) of attend-left valid hit condition relative to attend-right valid hit condition as a function of the number of electrodes (dimensions) used to calculate the projections. This analysis is done for electrode numbers for which at least 15 sessions were available, as indicated by the number on top of the error bar. The data point at the right end represents the projection calculated using all the electrodes of a session, which is also plotted in figure 5.

## Discussion

We show that nearly every measure of neural activity, including power spectral density, phase consistency, noise correlations and discriminability during neutral cueing is intermediate to those in the cued and uncued conditions, consistent with the effects observed earlier in spiking activity and behavior (Mayo and Maunsell, 2016). For single electrode analysis, the discriminability between attention conditions was higher for both gamma (42-78 Hz) and high-gamma (122-198 Hz) power compared to spiking. When multiple electrodes were used, population projections obtained using weights proportional to the difference in mean activity between correctly detected attend-left and attend-right conditions (as done in the “Attention Axis” analysis) were equally discriminable with spikes and high-gamma but poorer with gamma and alpha. However, once the full covariance matrix was incorporated using LDA analysis, the discriminability of high-frequency LFP power (both gamma and high-gamma) was comparable to spiking activity. These results were largely consistent across a wide range of attention conditions, including valid, invalid, and neutral cues.

### Relation to previous studies

LFPs have been previously used to decode stimulus information (Belitski et al., 2008; Kanth and Ray, 2020), saccade location (Pesaran et al., 2002), allocation of attention (Tremblay et al., 2015), motor movements (Bansal et al., 2012), and in brain-machine-interfacing (BMI) applications (Hwang and Andersen, 2013; Andersen et al., 2014). Consistent with our findings, many of these studies showed high decodability using high-frequency (>40 Hz) LFP. Similarly, BMI studies have shown that while single-channel LFPs offer better decoding of reach targets than single-channel spikes, decoders using multi-channel LFPs performed worse than multi-channel spikes because of larger noise correlations in LFPs than spikes, with comparable spike and LFP performance with ~8 electrodes (Hwang and Andersen, 2013).

A recent study analyzed neuronal responses in macaque primary visual cortex (V1) during attend-in, attend-out and attend-both (neutral) conditions to test whether spike-count correlations show a graded response as a function of the focus of attention (Denfield et al., 2018). They showed that the effect of attention on spike-count correlations depended on the duration over which analysis is performed. For short durations (100-200 ms), attention decreased spike-count correlations and neutral cueing produced intermediate correlations (see their Figure 5C). However, for longer analysis intervals, correlations for attend-in and attend-out were comparable, and attend-both produced the largest correlations. They suggested that the increase in correlation could be due to a switch in the focus of attention in the attend-both condition when observed over long time scales, which has also been reported in other studies (Landau and Fries, 2012; Landau, 2018; Fiebelkorn and Kastner, 2019). However, here we performed all analyses over 500 ms (mainly to improve the frequency resolution for spectral analysis of the LFP) and found that neutral cueing produced intermediate correlations when averaged over the two target locations (although, when separated based on the target location, correlations for target-in and target-out neutral hits were comparable to attend-out and attention-in valid hit conditions; Figure 4). We also found no evidence of attention switching, which would have led to negative spike-count correlations for interhemispheric pairs of electrodes. Instead, we found interhemispheric spike-count correlations to be small (0.044 ± 0.0012), consistent with previous reports where this analysis was done over short (200 ms) periods (Cohen and Maunsell, 2010; Mayo and Maunsell, 2016). Likewise, correlations in the LFP power at different frequencies were also not negative. Denfield and colleagues (2018) could not quantify the attentional switch using interhemispheric spike-count correlations because they did not record from both hemispheres simultaneously. It is possible that attentional switching is difficult to capture using inter-hemispheric spike-count correlations and may need more sensitive methods (Landau, 2018; Fiebelkorn and Kastner, 2019). The discrepancy between studies could also be due to differences in experimental design (for example, Denfield and colleagues had 100% validity and no catch trials in the non-neutral condition, in order to minimize attention switching) that could have encouraged different attention strategies on the part of their animals.

### Effect of attention on gamma oscillations

Previous studies have suggested a potentially important role of the phase synchronization of gamma oscillations in communication (Fries et al., 2001; Fries, 2015). Even when gamma oscillations are not strong enough to be visible as a distinct bump in the power spectral density, measures such as field-field and spike-field coherence often show such a bump, suggesting a potentially important role of phase synchronization (Fries et al., 2008). We found similar results in our data; while power increased over a wide range of frequencies between ~40-200 Hz and beyond with attention, the increase in phase synchronization was localized to a narrower range with a clear peak between 40-80 Hz. However, this does not necessarily imply the presence of narrow-band phase synchronization, because measures of phase synchronization could themselves be less sensitive at higher frequencies. For example, small offsets in time correspond to larger fluctuations in phase at higher frequencies (Ray, 2015). Indeed, in spite of negligible coherence, the discriminability of high-gamma (>122 Hz) was comparable (using LDA) or higher (other methods) than gamma (42-78 Hz) that showed higher phase-synchronization. Further, spikes were not locked to the phase of gamma rhythm but instead to the stimulus that was counterphasing at 10 Hz. Therefore, even though we observed increased field-field synchronization with attention in the gamma range, whether this synchronization has a functional role is unclear (Ray and Maunsell, 2015).

While correlations in power across electrodes showed similar trends as spiking activity during valid and neutral cueing, the trends were different during invalid cueing, especially for gamma oscillations (Figure 4). We interpret these results with caution since the number of invalid trials was smaller than valid and neutral trials. Nonetheless, the striking difference in the correlation trends for gamma versus other measures could provide clues about switching of attention in invalid trials. Attention has been divided into several ‘subcomponents’ such as orienting, target detection and vigilance, which may be carried out by distinct brain areas (for a review, see (Posner and Petersen, 1990)). Even orientation of attention involves disengagement from the present location, shifting, and finally redeployment to another location, which may involve the interaction of many brain areas (Posner and Petersen, 1990). Importantly, to detect an invalid target, a potentially large shift of attention may be required (see Figure 3 of Mayo and Maunsell, 2016). Since gamma oscillations have been linked to a variety of neural mechanisms such as normalization (Ray et al., 2013) and gain control (Ni et al., 2016) that could be involved in attentional switching, the modulation of gamma could be different from other measures in the invalid case. More detailed experiments in which various subcomponents of attention are manipulated independently are needed to test these predictions.

Correlations were also modulated differently for firing rates versus LFP power when comparing hits versus misses in some cases (Figure 4). These differences could be related to the variety of different reasons behind a miss, including focused attention at the incorrect location, a unique momentary distraction, or a reduction in global attention at both locations. Power at alpha and gamma frequency bands, which could be influenced by global synchronization (Buzsaki, 2006), could be modulated differently for these conditions.

### Spatial extent of selective attention

A surprising finding of this study is that discriminability of high-gamma remains comparable to spiking activity even when many electrodes are used (Figure 6), unlike other studies in which multi-channel spikes outperform multi-channel LFPs (Hwang and Andersen, 2013; Tremblay et al., 2015). This is reminiscent of the finding that electrocorticogram (ECoG), which have a much larger spatial spread than LFPs (Dubey and Ray, 2019), nonetheless show better decodability of natural images than LFPs or spikes (Kanth and Ray, 2020). Since spiking activity represents only a noisy estimate of the true underlying attentional state, averaging across a larger population of neurons (as done implicitly in population signals like the LFP) reduces noise and provides a more reliable estimate of the common signal (here, the attentional state), which likely explains why single-channel LFPs outperform single-channel spikes. The optimal level of neural integration for decodability of a particular stimulus or behavioral state depends on the spatial extent over which neural assemblies remain coherent under that stimulus/state. For example, if spatial attention is mediated through circuits involved in normalization, as suggested recently (Reynolds and Heeger, 2009; Verhoef and Maunsell, 2017), the spatial extent of normalization could determine the spatial resolution of attention. Since such divisive normalization mechanisms incorporate an inhibitory network that is involved in generating gamma oscillations (Carandini and Heeger, 2011; Ray et al., 2013), the spatial resolution of attention could be related to the spatial spread over which gamma oscillations are coherent. We and others have previously shown that the spatial spread of LFP is local (radius of ~0.5 mm; (Xing et al., 2009; Dubey and Ray, 2016)), with a slightly larger spread in the high-gamma range (Dubey and Ray, 2016, 2019). If this spread matches the spatial extent of selective attention (which could be dependent on the constraints of the task), decodability is likely to be best using high-gamma power. Our results therefore provide an indirect measure of the spatial scale over which attention might be operating.

## Conflict of interest

The authors declare no competing financial interests.

## Acknowledgements

This work was supported by Wellcome Trust/DBT India Alliance (Senior fellowship IA/S/18/2/504003 to SR) and DBT-IISc Partnership Programme and a National Institutes of Health Fellowship F32EY02259 to JPM. SSP, AD, STK were each supported by individual research fellowships in IISc, funded by the Ministry of Human Resource Development, Government of India. We thank Drs. Marlene Cohen and John Maunsell for helpful comments.

---

**Supplementary Figure 1:**
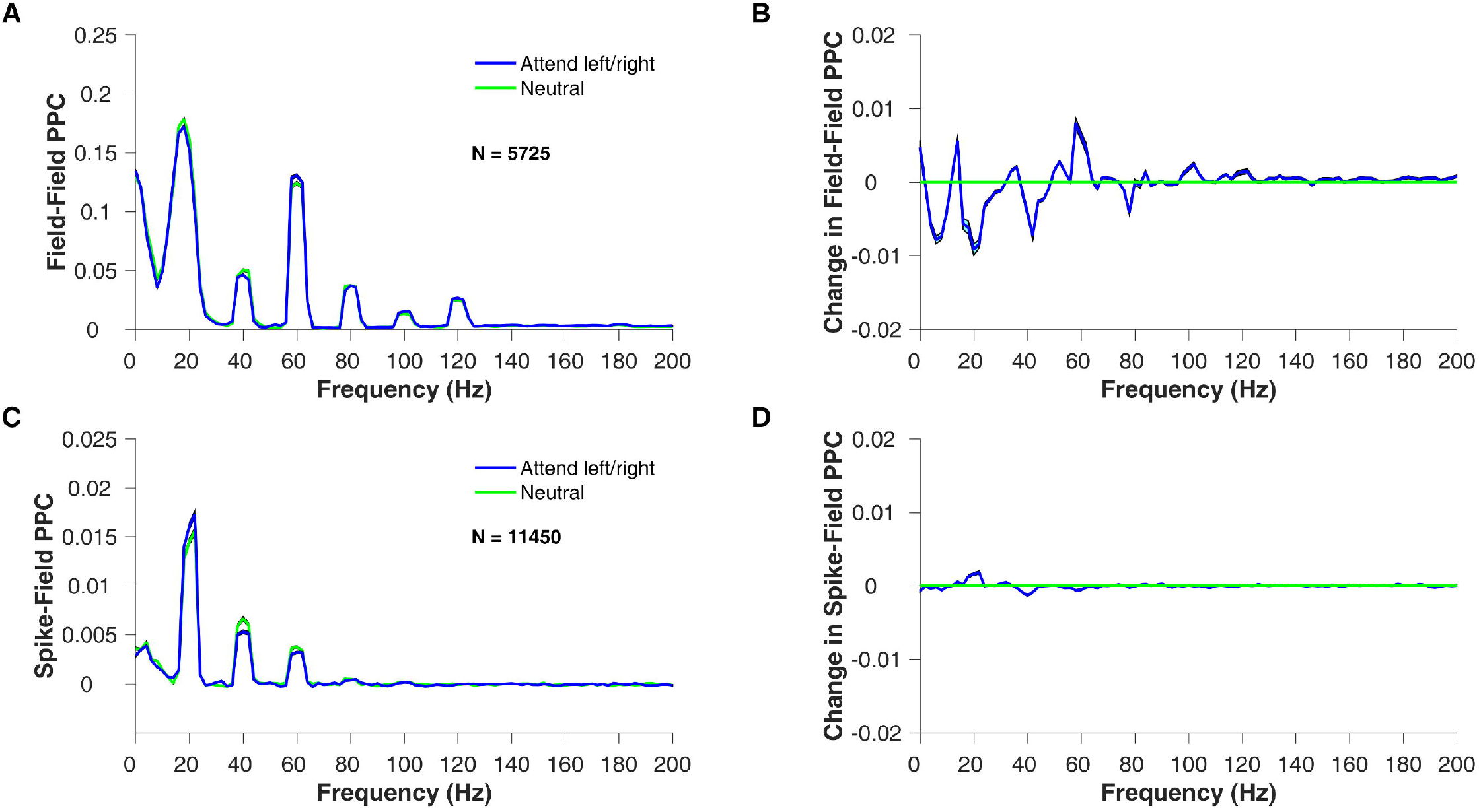
Inter-hemispheric electrode pairs phase relationship. **(A)** Mean field-field pairwise phase consistency (PPC) averaged across 5725 electrode pairs of 25 sessions of the attend-right/left (average of attend-left valid hit and attend-right valid hit condition; blue trace) and neutral (target-right neutral hit and target-in neutral hit combined; green trace) condition. **(B)** Mean change in field-field PPC of the attend-left/right condition (blue) relative to the neutral condition. **(C)** Mean Spike-field pairwise phase consistency (PPC) averaged across 11450 electrode pairs (including spike and LFP of the same electrode) for the same attention conditions as above. **(D)** Mean change in Spike-field PPC relative to the neutral condition.

**Supplementary Figure 2:**
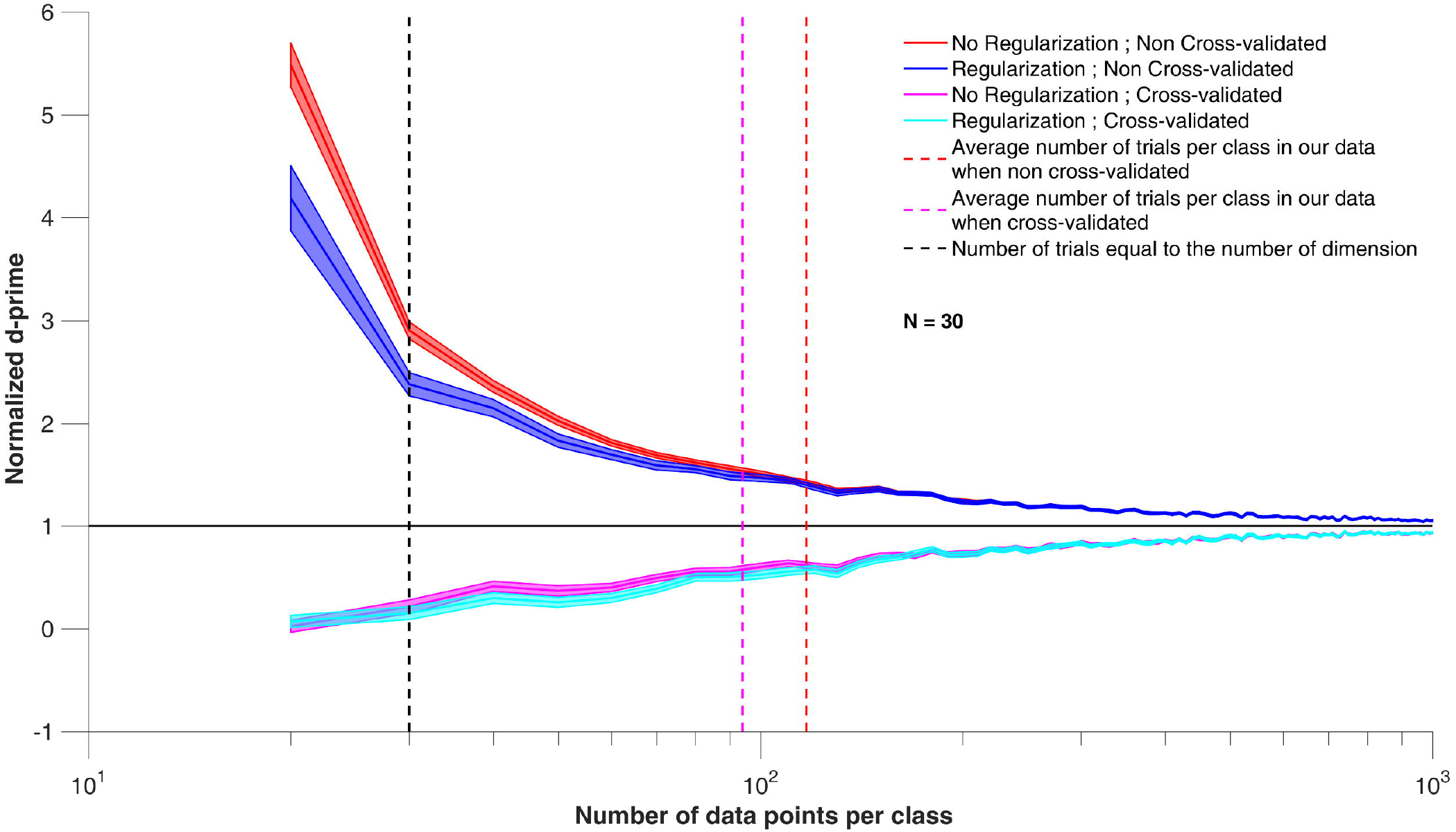
Effect of the number of data points per class/category on the estimation of d-prime between two populations using LDA and explicit regularization of LDA, with no cross-validation and 5-fold cross-validation. Estimated mean d-prime normalized relative to true d-prime between the projections of the populations as a function of the number of data points chosen per class to estimate the linear discriminant analysis (LDA) axis. The cases with no regularization with no cross-validation (red trace) and 5-fold cross-validation (magenta), regularized LDA with no cross-validation (blue) and 5-fold cross-validation (cyan) are compared. N (=30) is the number of dimensions used, which is close to the average number of electrodes per session in our dataset. The black vertical dashed line indicates the case when the data points are equal to the number of dimensions. The magenta and red vertical dashed lines indicate the average number of data points in our dataset with and without cross-validation, respectively.

**Supplementary Figure 3:**
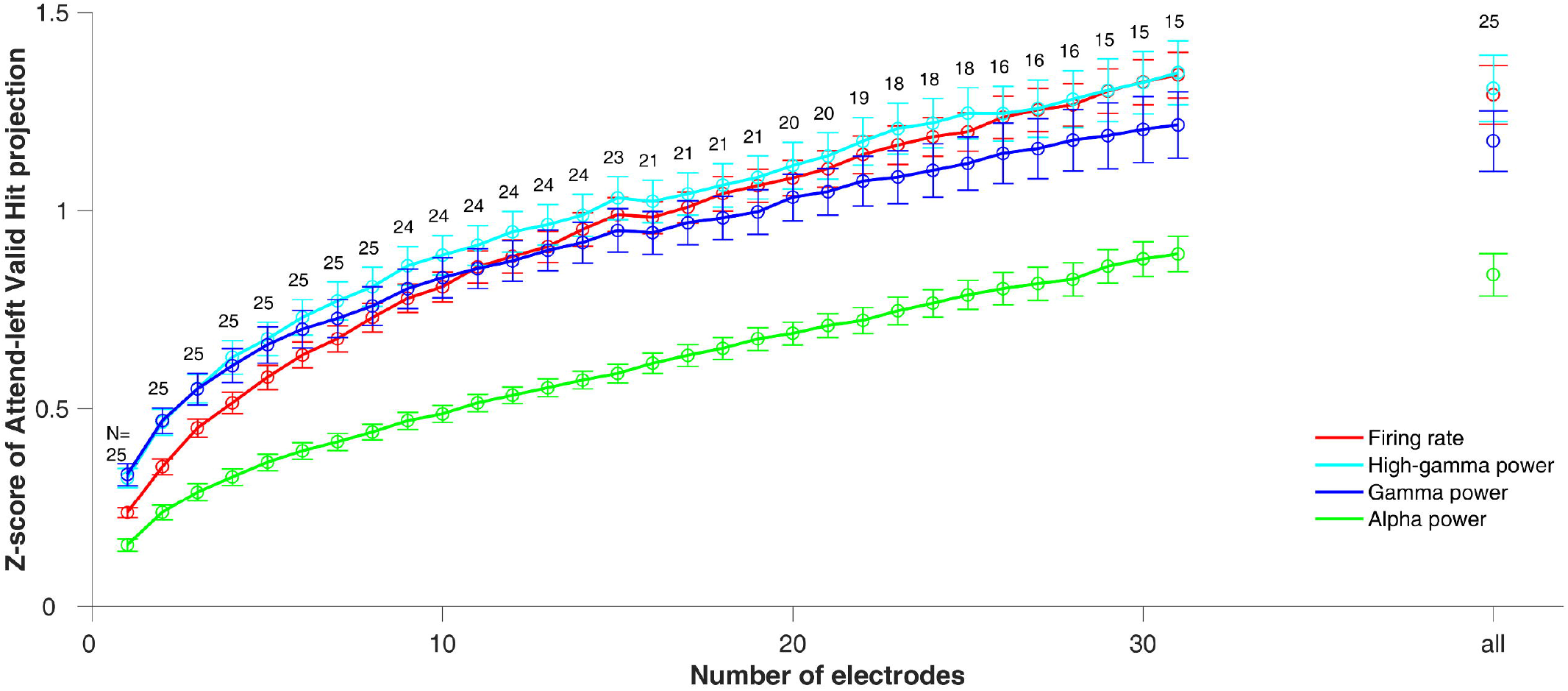
Discriminability as a function of the number of electrodes without cross-validation. Same as Figure 6 but without any cross-validation. Note that unlike the cross-validated case, values do not asymptote here, likely due to an increase in the degree of over-fitting for larger dimensionality.

## Notes

### Competing Interest Statement

The authors have declared no competing interest.

